# Clinical diagnosis of single-gene disorders and chromosomal abnormalities based on BGISEQ-500 platform

**DOI:** 10.1101/675991

**Authors:** Yanqiu Liu, Liangwei Mao, Xiaoming Wei, Jianfen Man, Wenqian Zhang, Lina Wang, Long Li, Yan Sun, Wei Li, Teng Zhai, Xueqin Guo, Lique Du, Jin Huang, Hao Li, Yang Wan, Hui Huang

## Abstract

Most of the variation in the human genome is a single nucleotide variation (SNV) based on a single base or small fragment insertions and deletions and genomic copy number variation (CNV). Both types of mutations are involved in many human diseases. Such diseases often have complex clinical symptoms and difficult clinical diagnosis, so an effective detection method is needed to help clinical diagnosis and prevent birth defects. With the development of sequencing technology, the method of chip capture combined with high-throughput sequencing has been extensively used because of its high throughput, high accuracy, high speed and low cost. This study designed a chip that captures the coding region of 3043 genes associated with 4013 monogenic diseases. In addition, 148 chromosomal abnormalities can be identified by setting targets in specific regions. Compared with the whole exon chip, the chip can detect 4013 monogenic diseases and 148 chromosomal abnormalities at a lower cost, including SNV, intra-gene CNV and genomic copy number variation. This study utilized a strategy of combining the BGISEQ500 sequencing platform with the chip to identify 102 disease-associated mutations in 63 patients, 69 of which were new mutations. The evaluation test results also show that this combination complies with the requirements of clinical testing and has good clinical application value.

## Introduction

Monogenic diseases involve multiple disciplines, with complex clinical symptoms and difficult diagnosis, and most of them are usually fatal, disabling, or teratogenic^1^. Traditional testing techniques have the risk of a missed diagnosis and misdiagnosis, which may lead patients to miss the best treatment opportunity. Genetic testing can achieve early detection, early intervention, and early treatment of single-gene genetic diseases. The consequences of large-scale gene discovery and validation of monogenic diseases can be quickly promoted and applied clinically. People with a family history of genetics can detect and avoid the birth of children with birth defects by pre-marital, pre-pregnancy and prenatal genetic screening ^2,3^ Therefore, genetic testing is important for clinical diagnosis and prevention of birth defects.

Next-generation sequencing technology has been widely used in genetic disease detection. The main technical solutions are targeted region sequencing, whole exome sequencing, whole genome sequencing and mitochondrial DNA sequencing. The biggest advantages of high-throughput sequencing technology are low cost and high throughput ^4^, which are commonly used in genetic disease detection and carrier screening ^5^. Whole genome and exome detection are not only costly and time consuming, but screening for disease-causing mutations in so much data is also a huge challenge ^6^ The combination of regional capture and high-throughput sequencing technology can effectively capture disease-associated regions, quickly find disease-causing mutations, and has the characteristics of high throughput, low cost, high speed, and high accuracy, and is widely used in clinical practice ^7,8^. However, most of the current genetic testing products detect fewer types of diseases and have a lower detection rate. In addition, diseases such as mental retardation and growth retardation, in addition to single-gene genetic diseases, recent studies have found that chromosome microdeletions or microduplications are also important causes of developmental delay and mental retardation^9^. Therefore, we urgently need a highly efficient, sensitive detection method that can be found for all types of mutations to meet the needs of one-step detection of a variety of monogenic genetic diseases and common chromosomal abnormalities.

Therefore, this study used BGISEQ-500 as a sequencing platform to develop a chip that focuses on coding regions with known associations with genetic diseases. Variants that affect gene function are detected more cost-effectively than whole genome or whole exome. Currently, 4013 known single genetic diseases can be detected (Table 1). In addition, we can detect 148 common chromosomal abnormalities by targeting specific regions (Table 2). Compared with traditional gene detection methods, we integrate known single gene diseases and common chromosomal abnormalities, and achieve “one-stop” solution to genetic mutations, which can improve the detection rate of diseases, with the advantages of high throughput, high accuracy, fast speed and low cost. It is a powerful tool for clinical diagnosis and prevention of birth defects.

## MATERIALS AND METHODS

### Sample information

A total of 100 samples were gathered for this study. In order to assess the stability of the chip, we selected two samples S77 and S78 for inter-batch and intra-batch stability evaluation. In addition, samples S79, S80, S81 and S82 were selected to evaluate the coverage and depth of the target area under the BGIS EQ500 platform. 86 patients were selected from clinical cases. Among them, 63 cases were diagnosed and 23 cases were not. In addition, 12 samples that have been tested for CNVseq were selected, and the results were in accordance with the known positive samples in the disease area shown in Table 2. The ability of the chip to detect chromosomal abnormalities was evaluated. All adult participants and parents of minors registered in the study have obtained written informed consent. The project and research programmes involving human tissues were approved by the BGI-shenzhen (BGI-IRB 16098) Ethics Committee.

### Chip design

In this study, a chip was designed to detect not only SNP, INDEL and large intragenic deletion, but also 148 chromosomal abnormalities from DECIPHER and OMIM databases by adding capture fragments in specific regions. The design steps of capture region are as follows: (I) Design of capture region of single gene disease: Because a gene often corresponds to multiple transcripts, first, select one most commonly used or longest transcript for each gene as the only transcript corresponding to the gene, then select all CDS regions, each CDS region extending 10 bps on both sides, to detect splicing variation. The UTR region is large, and most of the regions cannot be annotated. The chip does not capture the UTR region. The transcripts selected according to the above principles may not include all the functional regions of the other transcripts of the gene, and finally complement the missing functional regions; (II) Design of chromosomal abnormality detection capture region: Firstly, the mutation region of each chromosomal abnormality and all genes in the region are determined according to the database, and then one of the most commonly used or longest transcripts is selected for each gene. All functional regions of the major pathogenic genes of the mutated region are all intended to be contained within the chip. The capture regions of other non-major pathogenic genes are based on the following principles: (i) For mutation regions less than or equal to 15 genes, we randomly select 100 bps CDS region of each gene for capture; (ii) For mutation regions with more than 15 genes and less than 40 genes, CDS 100 bps of two genes was randomly selected for capture for each three genes; (iii) For mutation regions larger than or equal to 40 genes, CDS 100 bp was randomly selected for each gene interval.

### Experiments and sequencing

In this experiment, genomic DNA was first extracted from whole blood, and qualified DNA was subjected to library preparation(7). The library was prepared by disrupting 1 μg of genomic DNA into a small fragment of 200-300 bps of DNA. The fragment selection product was quantified using Qubit. The initial amount of DNA was adjusted to 50 ng according to the measured concentration. The end repair is then performed and the base “A” is ligated at the 3’ end so that the DNA fragment can be ligated to Barcode. The library constructed by Pre-PCR was used to enrich the target region by the probe designed in this study. The enriched product was amplified by PCR. Finally, qPCR was utilized to detect the product before and after hybridization to obtain hybridization efficiency. After the qPCR assay passed, 160 ng of the DNA library was subjected to cyclization, and then DNA nanospheres were synthesized. Sequencing was performed using the BGISEQ-500 platform, and data analysis and interpretation of the results were performed based on the sequencing data^10,11^. The data that support the findings of this study have been deposited in the CNSA (https://db.cngb.org/cnsa/) of CNGBdb with accession code CNP0000378.

### Bioinformatics analysis and mutation identification

The process of bioinformatics analysis includes data filtering, alignment, mutation detection and result annotation. The raw data is first evaluated for quality to remove low quality and adapter contaminated reads. The valid data is then mapped to the human reference genome (HG19) using Burrows Wheeler Aligner (BWA) ^12^. The PCR-induced duplication was eliminated using Picard software. SNVs and Indels were tested using the Genomic Analysis Toolkit (GATK). Intra-gene deletions and duplications were identified by comparing the average depth between samples in the same batch. The mutations are then annotated, using databases including dblocal (a database of mutation frequencies for 100 normal human samples), dbSNP (http://www.ncbi.nlm.nih.gov/SNP/), HapMap (http://hapmap.ncbi.nlm.nih.gov/), dbNSFP (http://varianttools.sourceforge.net/Annotation/DbNSFP) and 1000 Genomes (http://www.1000genomes.org/). In addition, we used CNVkit to detect chromosomal abnormalities ^13^. Finally, suspicious mutations are screened, interpreted and validated to generate the final result. The bioinformation analysis method of CNVseq refers to ^14^ The clinical evaluation of CNVseq results is based on guidelines prepared by the American College of Medical Genetics (ACMG) ^15^. Mutations are named in reference to the International Cytogenetic Nomenclature International System (ISCN) standard.

### Homology model construction and protein stability prediction

To investigate the effects of disease-associated mutations on protein structure and function, we performed protein modeling analysis. This study used the rapid modeling module of yasara (version 17.1.28) software to automatically implement multi-template search and comparison to complete hybrid modeling. The protein model was repaired using the foldx plug-in and the mutated protein model was constructed to obtain a high confidence structural model. The foldx plug-in was used to calculate the difference between the energy of the wild-type protein and mutant protein (ΔΔG = Δgmut-Δgwt), and a value of ΔΔG greater than 1.6 kcal/mol was considered to have a significant effect on protein stability ^16^. All structural analysis and image rendering were performed using PyMOL (version 2.2.0).

## Results

### Clinical sample depth determination

In this study, four samples S79, S80, S81 and S82 were selected. The original depths of the samples were 281.46X, 413.23X, 569.57X and 714.32X, respectively. The depths after removing the duplication were 157X, 231X, 277X and 380X, respectively. All CDS regions were screened by parameters with depth greater than or equal to 30X and coverage greater than or equal to 95%. The coverage of CDS regions of four samples with different sequencing depths were S1 95.64%, S2 97.73%, S3 98.06% and S4 98.80% (Fig. 1). Based on the above results, it is recommended that on the BGISEQ500 sequencing platform, the depth of sequencing of the samples using the customized chip of this study should preferably reach 200X or more after the removal of the duplication.

**Figure 1.**
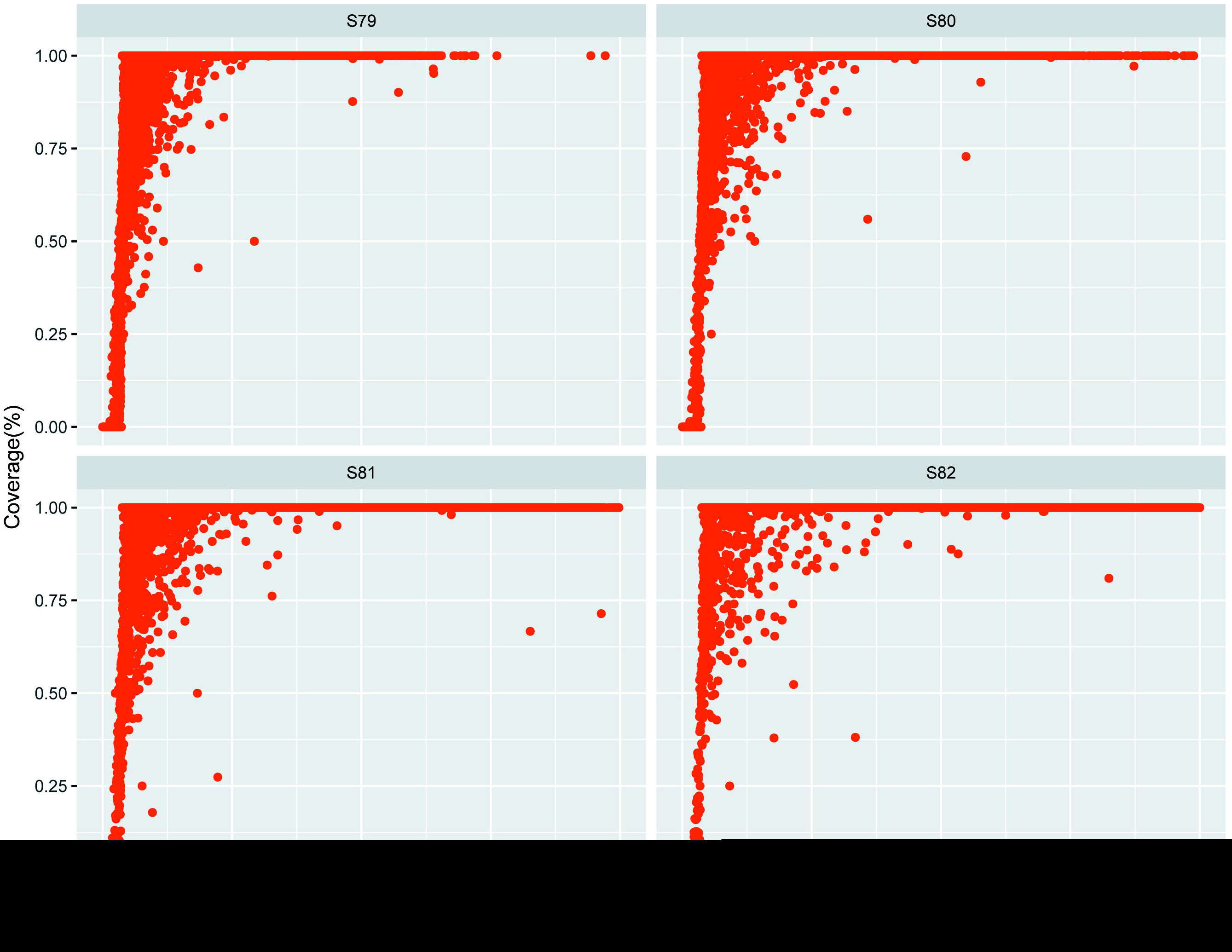
Relationship between Sequencing depth (more than 30×) and coverage (more than 95%) in CDS region

### Inter-batch and intra-batch stability assessment

In this project, sample S77 and sample S78 were sequenced in three batches to evaluate the stability among batches; each sample was sequenced three times to evaluate the stability within the batch. We use the parameter - out_mode EMIT_ALL_SITES to output all locus detection information in the capture region. Genotype consistency of loci in different batches of the same sample and the same batch of repeated samples was analyzed. For batch-to-batch stability, the total number of loci was 9903792 for sample S77, the intersection of three different batches was 9881645, the stability was 99.78% Fig. 2(a); 9874160 for sample S78, 9852762 for three separate batches, 99.78% for stability Fig. 2(b). For intra-batch stability, the total number of loci in sample S77 was 9904450, the intersection loci of three samples in the same batch were 9882 238, the stability was 99.78% Fig. 2(c); sample S78, the total number of loci was 9877 841, and the intersection loci of three samples in the same batch were 9857 175, the stability was 99.79% Fig. 2(d). From the above data, it is confirmed that the stability of the customized chip is quite good among batches and within batches on the BGISEQ500 sequencing platform.

**Figure 2.**
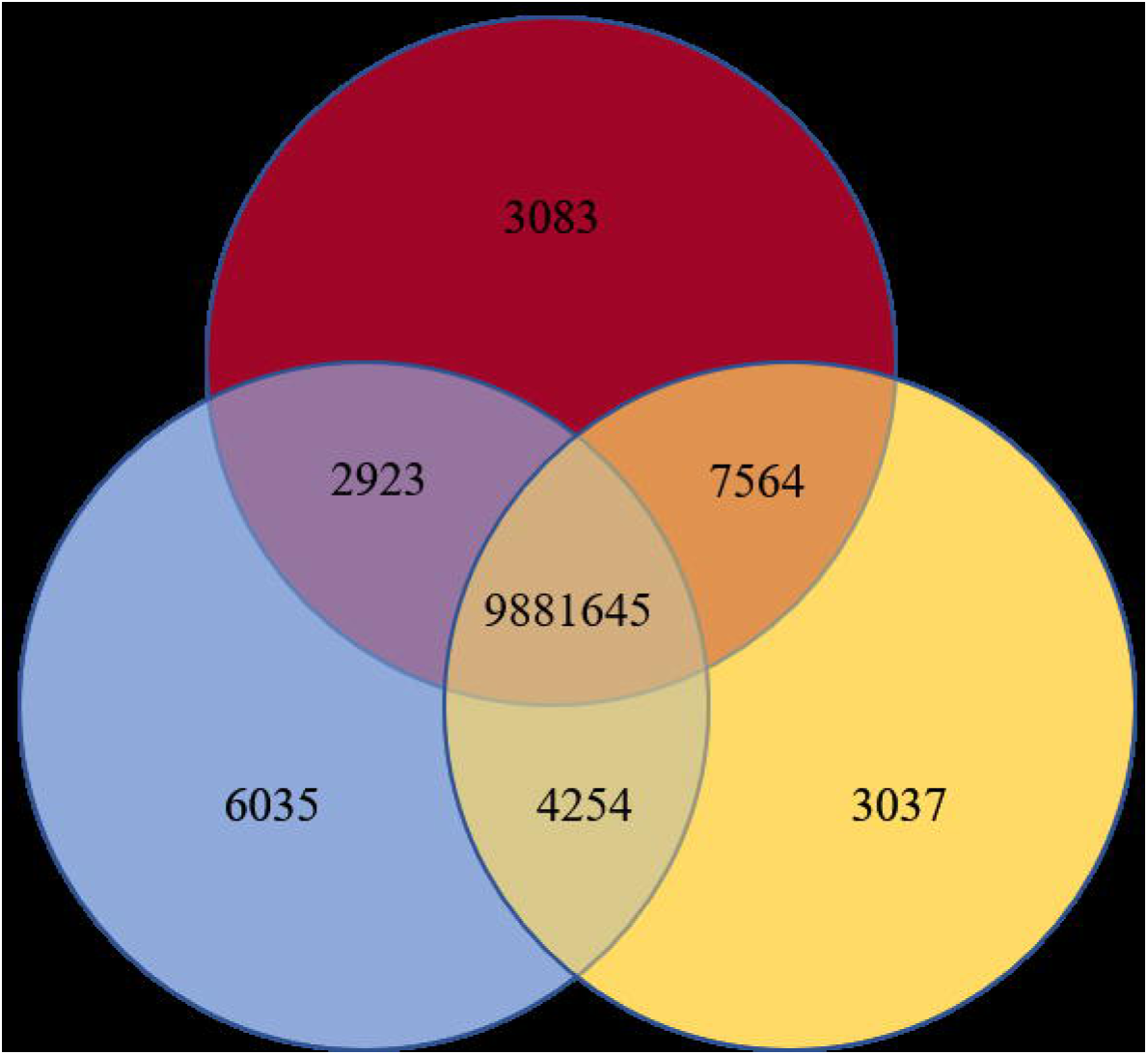
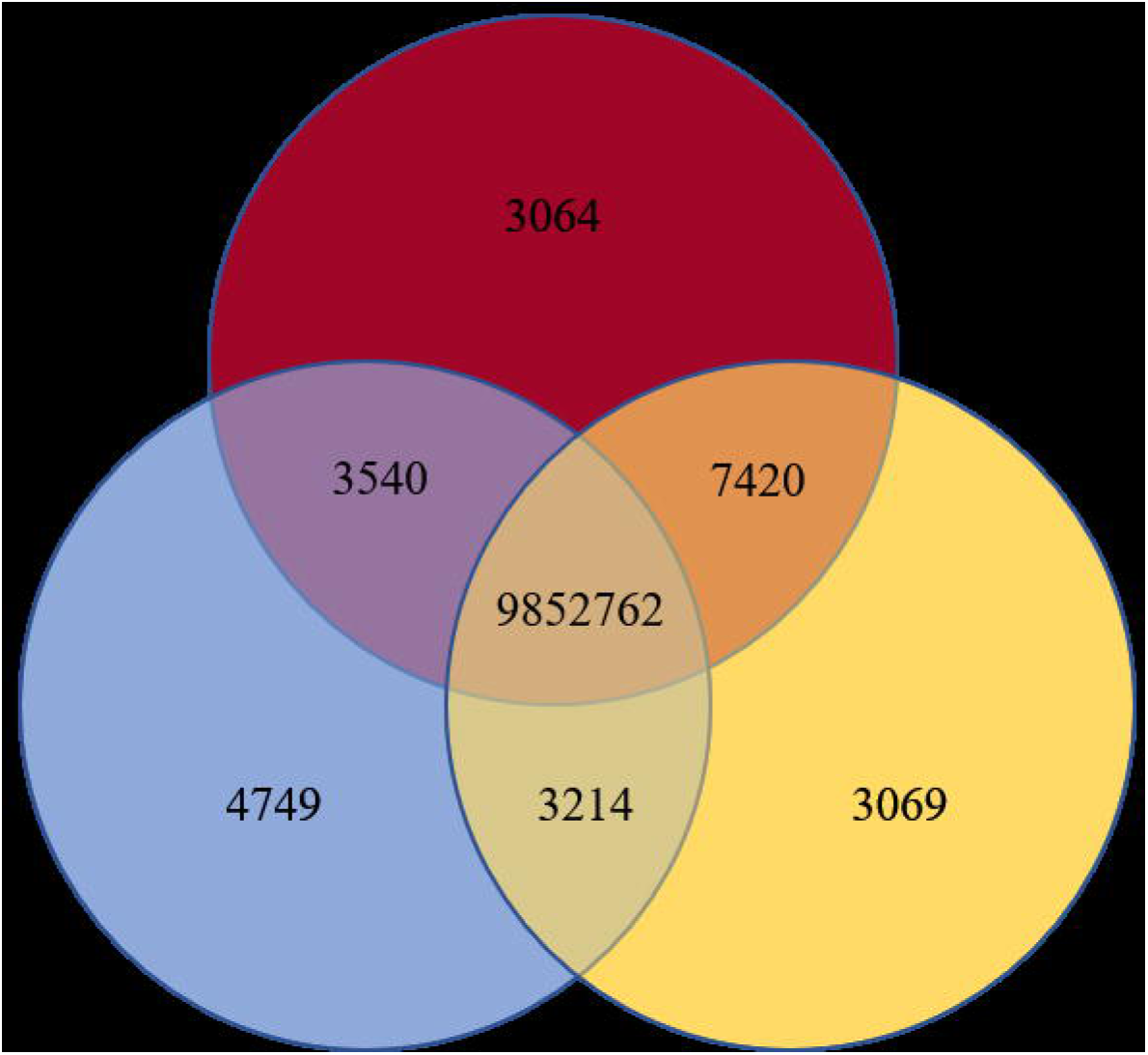
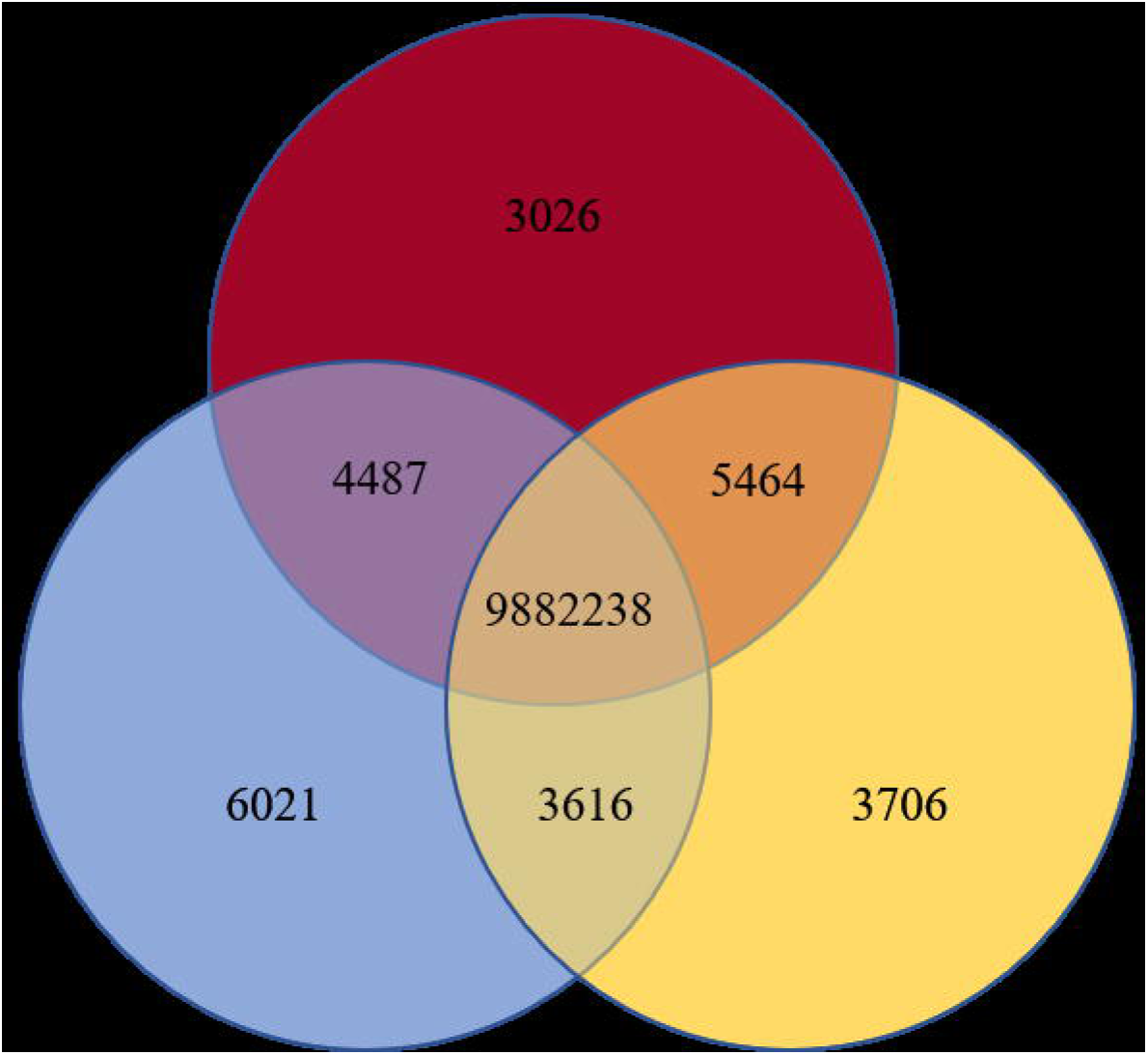
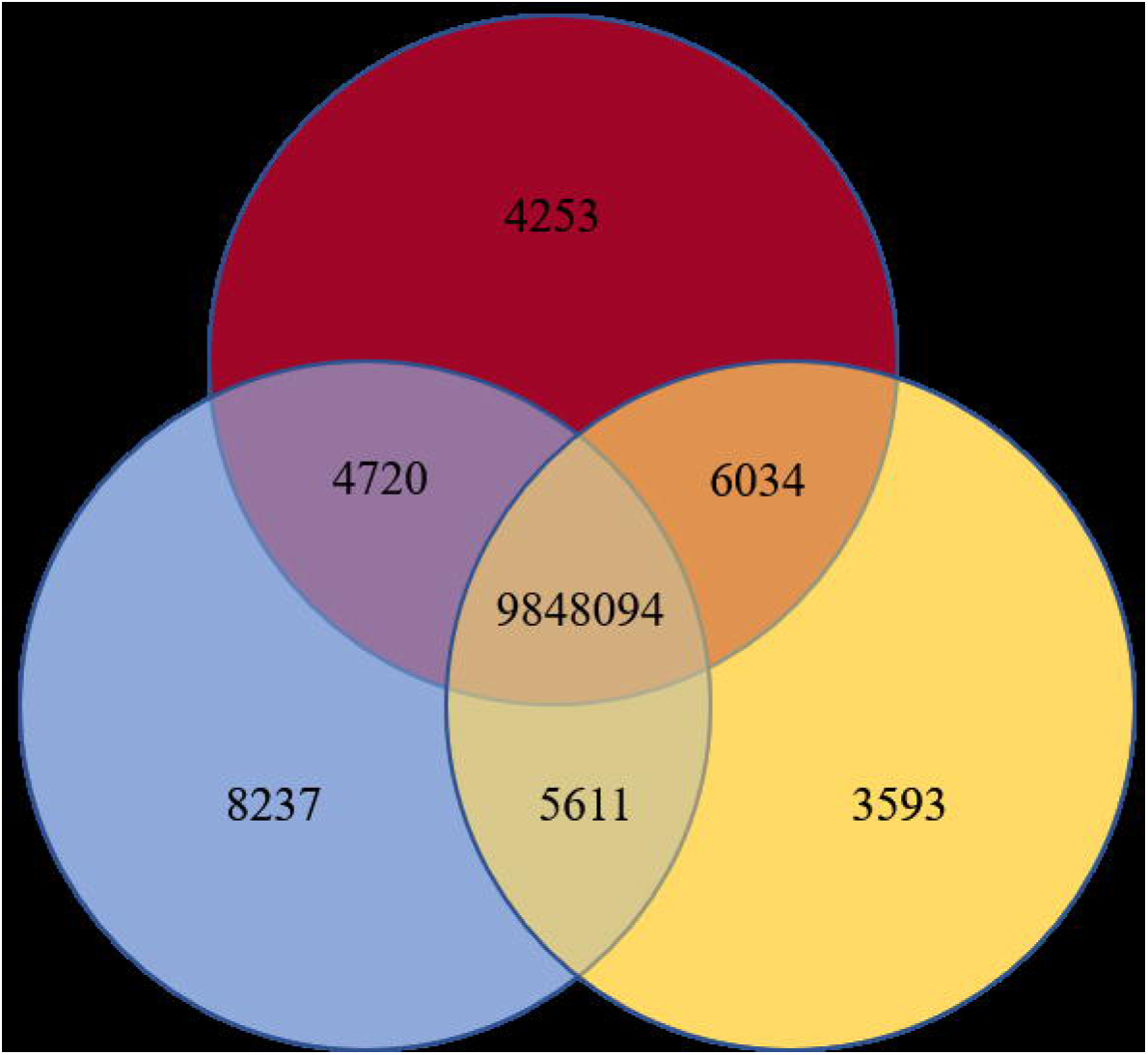
Evaluation of the stability of our method. (a) Venn diagram of S77 sequenced three times in a same batch. (b) Venn diagram of S78 sequenced three times in a same batch. (c) Venn diagram of S77 sequenced three times in a same batch. (d) Venn diagram of S78 sequenced three times in a same batch.

### Mutation information in clinical samples

Using targeted NGS, we obtained high quality sequences of 86 samples. Mutation-related information is obtained after the completion of the reference sequence alignment and mutation detection. In this study, 102 disease-related mutations were identified in 63 patients, including 80 missense mutations, 8 frameshift mutations, 5 splicing mutations, 5 nonsense mutations, 3 intra-gene deletions, and 2 whole gene deletions. Of all the samples tested, 21 sample family information was valid. After verification of pedigree cosegregation, there are two de novo mutations among these families. Of the 102 mutations, 33 have been reported and 69 have been reported for the first time. Table 3 summarizes the disease-related mutation information for 63 samples.

### Chromosome abnormality detection

This study used CNVkit software to detect chromosomal abnormalities. The software detects CNV based on the read depth method. Therefore, in addition to the original depth, 10000000 reads and 20000000 reads are randomly extracted, simulating different sequencing depths for CNV copy number and breakpoint position detection. When the data shows that the original depth is 613 ×, there is still one area that is not detected. This area is: chr7: 69, 783, 279-69, 952, 448, the segment length is: 169.17 Kb, and the area is not detected at three different depths, namely 613 × (original depth), 140 × (20000000 reads) and 70 × (10000000 reads). Therefore, it is speculated that the detection accuracy of the customized chip is insufficient to detect a deletion and duplication of about 200 kb. In addition, the recommended detection accuracy of CNVkit software is 1M, and it is found that all the deletions and repetitions above 1M are detected. CNVkit software detects chromosome deletions and duplications based on depth of reads. The results also confirmed that as the depth decreases, the number of missed detection areas increases, so it is recommended to ensure a certain amount of depth to help reduce the rate of missed detection. Table 4 shows the sample detection information.

### Protein structure prediction and stability results

We performed protein modeling analysis on all genes defined as uncertain significance, of which only seven genes were modeled completely and the sequences included mutant amino acids (Table 5). The seven genes are: *ANLN, SLC7A9, CNGB1, UMOD, DSTYK, UNC45B, COL4A3*. In the structure of ANLN, Asp1021 is located at the carboxy terminus of the Anillin protein and belongs to the PH (Pleckstrin homology) domain, which is necessary for all targeted events ^17^. The PH domain is a 120 amino acid protein module that is thought to interact with lipids to mediate protein recruitment to the plasma membrane, and studies have shown that the PH domain is electrostatically polarized ^18^. To examine how the p.D1021V mutation would affect protein structure, we compared the structure of wild-type and mutant, and found that the conformation is basically unchanged. In addition, Gibson free energy calculated by foldx also indicates that the mutation does not affect the stability of the protein.

The protein b0, +AT (encoded by the *SLC7A9* gene) and protein rbAT (encoded by the *SLC3A1* gene) form a heterodimeric amino acid transporter through disulfide bonds, characterized as a Na+-independent transport system mediating obligatory exchange of various cations and neutral amino acids including cystine. The protein b0, +AT is a typical helix globin with 12 transmembrane helices presumed by hydrophilicity analysis ^19^. The mutation p.T77M is located in the second transmembrane helix(fig3(a)). From the results of structural and Gibbs free energy changes, we have not found any cause that can affect protein function. However, the p. T123M mutation is deemeded to be pathogenic, resulting in a moderate phenotype^20,21^. p.T123M is located in the third transmembrane region and is in the middle of the transmembrane helix (fig3(b)) as the mutant p.T77M. Although it is impossible to infer how these two mutations affect protein function, we speculate that they are the same type of mutation.

**Figure 3.**
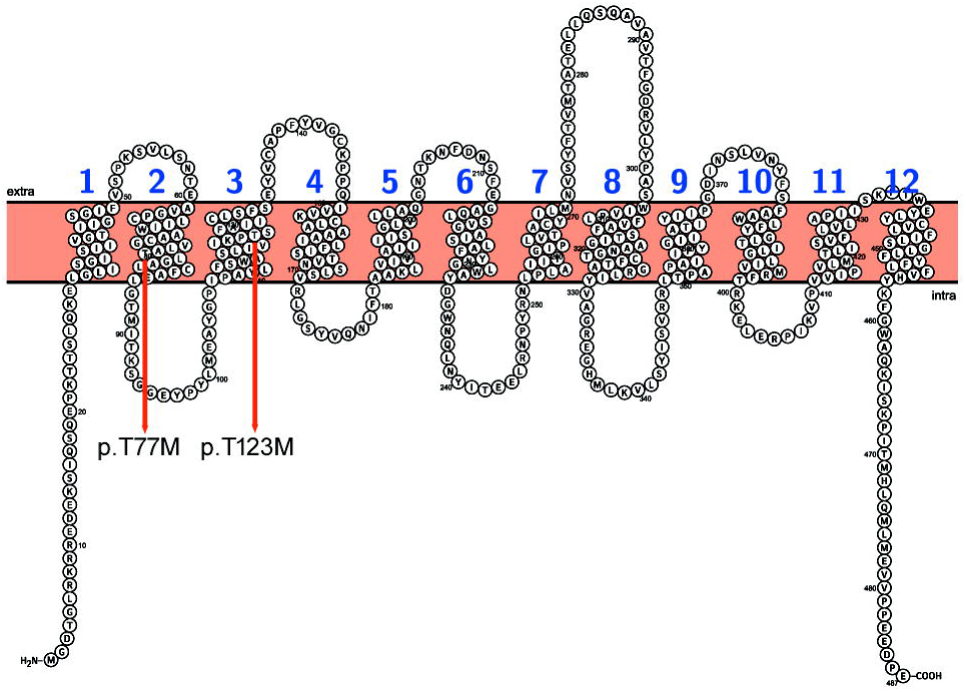
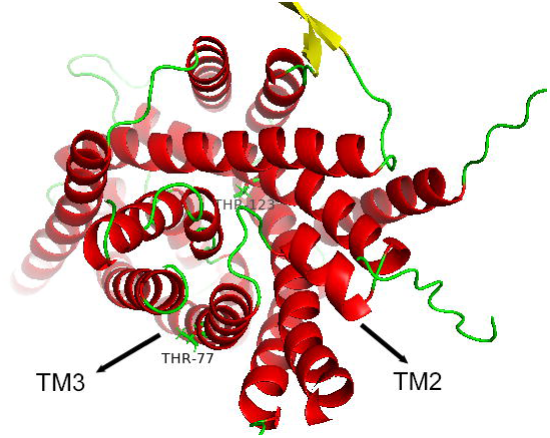
The SLC7A9 protein’s topolog and 3D structure. (a) Illustration of SLC7A9 protein’s topolog. The red arrow is marked as the location of the mutation. (b) The structure of the SLC7A9 protein is shown using cartoon feature in PyMOL. The helices were shown in red colour, the β stands in yellow and the loop in green color. Further, the transmembrane (TM) helices number were labeled to better understand the location of the mutations in the protein structure.

The CNGB1 mutation p.M974R, UMOD mutation p.V550I, UNC45B mutation p.C786Y and COL4A3 mutation p.A1555V were calculated by foldx, the change ΔΔG in Gibbs free energy, to be 4.07063 kcal/mol, 4.01864 kcal/mol, 3.16846 kcal/mol and 2.46126 kcal/mol, respectively. This indicates that these mutations affect the stability of the protein. The DSTYK mutation p.R592Q change from large size and basic to medium size and polar, but we still have not found evidence of the mutation causing disease from structural changes or stability calculations.

## Discussion

The study of monogenic hereditary diseases belongs to the field of typical precision medicine. The complex clinical symptoms of monogenic diseases lead to difficult diagnosis, and most of the pathogenic mechanisms are not clear. Due to the lack of effective treatments, the disease is often fatal, disabling or teratogenic. Diseases such as mental retardation and growth retardation are often caused by chromosomal abnormalities in addition to the causes of monogenic genetic diseases. Therefore, we urgently need an effective detection method to help clinical diagnosis and prevention of birth defects, which can detect both monogenic genetic diseases and chromosome aberrations. This study designed a chip that can detect up to 4,013 single-gene diseases. In addition, this study also identified 148 common chromosomal disorders by setting targets on key genes and randomly setting targets at non-critical genes in chromosomal abnormal regions. Compared to whole exome sequencing, this custom capture chip enables the detection of various types of mutations in the sample at a lower cost. This project uses the strategy of BGISEQ500 sequencing platform and chip combination. The evaluation results show that this combination meets the requirements of clinical testing and has useful clinical application value.

For 86 clinical cases, we first found candidate pathogenic genes in the list of 4013 diseases based on clinical diagnosis, and used the targeted NGS to find pathogenic mutations in the candidate genes. If the mutation is indeterminate based on the results of the information analysis and database annotations, we will plot the reads and align the reference sequences of the mutation sites with a single base resolution. If the mutation is still unrecognized, Sanger sequencing or real-time PCR will be performed. However, the pathogenic mutations in some cases are still not in the candidate gene. We will find candidate mutations in other genes in the target region and to infer the disease in reverse.

In this study, we performed homology modeling on some proteins, hoping to be able to explain the changes in protein structure from mutations. Sample S32, 7 years old, clinical manifestations of hematuria and C3 glomerulopathy. Missense variation c.3062A>T (p.D1021V) was detected in the *ANLN* (NM_018685.4) gene coding region of the sample as a heterozygote. ANLN gene mutation can cause focal segmental glomerulosclerosis type 8 (OMIM#: 616032), which is autosomal dominant, the main clinical manifestation of glomerular segmental sclerosis, proteinuria, decreased glomerular filtration rate and progressive decline in renal function. Both SIFT and PolyPhe-2 predictions are deleterious mutations. The frequency information of c.3062A>T was not found in the dbSNP database, Hapmap database, thousand-person database and local database, and there is no documented pathogenicity. In the structure of ANLN, the mutation p.D1021V is located in the PH (Pleckstrin homology) domain. Anillin is an actin binding protein involved in cytokinesis. It interacts with GTP-bound Rho proteins and results in the inhibition of their GTPase activity. The PH domain has multiple functions, but generally involves targeting the protein to an appropriate cellular location or interacting with a binding partner. The PH domain is in electrostatic polarity, aspartic acid is charged and polar, and is often involved in the formation of protein active sites or binding sites, but proline is a non-polar amino acid. Comparing the wild-type and mutant conformations, no changes were found, but there were some differences in the hydrophobic surface. We speculated that the mutation affected the electrostatic polarity of the PH domain, resulting in a change in protein function. Therefore, it is speculated that the *ANLN* gene c.3062A>T mutation is a disease-causing mutation in the subject. Sample S36, 5 and a half years old, clinical manifestations of frequent urination, oliguria, hematuria. Missense variation c.230C>T (p.T77M) was detected in the *SLC7A9* (NM_014270.4) gene coding region of the sample, which was heterozygous. *SLC7A9* gene mutation can cause cystineuria (OMIM#: 220100), which is autosomal dominant or autosomal recessive. The main clinical manifestations are renal insufficiency, cystinuria, repeated urinary tract infection, Kidney stones and so on. The incidence of this mutation in the population is extremely low. There is not any literature on the pathogenicity of the *SLC7A9* c.230C>T variant. It has been previously reported in the literature that the p.T123M mutation is pathogenic and causes a moderate phenotype. This mutation p.T77M is in the middle of the transmembrane helix as the p.T123M mutation. Although it is not possible to infer how these two mutations affect protein function, we speculate that they are the same type of mutation. Therefore, it is speculated that the c.230C>T mutation of the *SLC7A9* gene is a disease-causing mutation in the subject.

CNV is widely distributed in the human genome and is one of the important pathogenic factors of human diseases. Pathogenic CNV can cause mental retardation, growth retardation, autism, various birth defects, leukemias and tumors. Determining the copy number and breakpoint position of the variant region is two crucial aspects of CNV detection. With the advancement of technology, more and more technical means for CNV detection, but different technology platforms and their corresponding computing strategies have great differences in the accuracy of CNV copy number and breakpoint position. The NGSseq method uses genome-wide data. This study utilizes genomic target region data. Although two methods for detecting CNV are based on the circular binary segmentation algorithm, there are still differences in data correction and comparison. Based on the above reasons, the position of the breakpoints obtained by the two methods is not very consistent. The breakpoint obtained by the NGSseq method is standard, and the difference between the breakpoint position and the standard position obtained by the chip is at the kb level. This study uses CNVkit software, which detects CNV based on the read depth method. Therefore, in addition to using the original data, we also simulated different sequencing depths for CNV copy number and breakpoint position detection. As the depth decreases, the number of missed detection areas increases, and a certain number of read lengths help to reduce the rate of missed detection. On breakpoint locations, different depths have no effect on the detection of breakpoint locations.

In summary, we provide a diagnostic detection tool that combines capture arrays and NGS to capture the coding region of 3043 genes associated with 4013 diseases, and detects 148 chromosomal abnormalities by setting targets in specific regions. The results of the evaluation indicate that our method has high accuracy and stability in detecting pathogenic mutations. Compared with traditional genetic testing methods, it integrates known single-gene diseases and frequent chromosomal abnormalities to achieve a “one-stop” solution to genetic mutations. This technology can be utilizes to diagnostic testing, providing an effective basis for clinical diagnosis and genetic counseling, and improving the detection rate of diseases.

## Acknowledgments

We gratefully thank all blood donors for their invaluable contribution to this study. Thanks to Dr. Zhongji Pu for helping with modeling and analysis of proteins.

## Author Contributions

Y.Q.L. and L.W.M. designed the research and wrote the first draft of the article. Y.Q.L. and Y.W. provided patient specimens and collected clinical information. X.M.W., L.L., Y.S., W.L., J.H., and H.L. designed and performed the experiments. Y.W., and H.H. contributed to drafting and revising the manuscript. J.F.M., W.Q.Z., L.N.W., T.Z., X.Q.G., and L.Q.D. performed data analysis. All authors reviewed the manuscript.

